# Real-time fMRI neurofeedback modulates auditory cortex activity and connectivity in schizophrenia patients with auditory hallucinations: A controlled study

**DOI:** 10.1101/2025.01.13.632809

**Authors:** Clemens C.C. Bauer, Jiahe Zhang, Francesca Morfini, Oliver Hinds, Paul Wighton, Yoonji Lee, Lena Stone, Angelina Awad, Kana Okano, Melissa Hwang, Jude Hammoud, Paul Nestor, Susan Whitfield-Gabrieli, Ann K. Shinn, Margaret A. Niznikiewicz

## Abstract

**Background and Hypothesis:** We have reported previously a reduction in superior temporal gyrus (STG) activation and in auditory verbal hallucinations (AHs) after real-time fMRI neurofeedback (NFB) in schizophrenia patients with AHs.

**Study Design:** With this randomized, participant-blinded, sham-controlled trial, we expanded our previous results. Specifically, we examined neurofeedback effects from the STG, an area associated with auditory hallucinations. The effects were compared to Sham-NFB from the motor cortex, a region unrelated to hallucinations. Twenty-three adults with schizophrenia or schizoaffective disorder and frequent medication-resistant hallucinations performed mindfulness meditation to ignore pre-recorded stranger’s voices while receiving neurofeedback either from the STG (n=10, Real-NFB) or motor cortex (n=13 Sham-NFB). Individuals randomized to Sham-NFB received Real-NFB in a subsequent visit, providing a within-subject ’Real-after-Sham-NFB’ comparison.

**Study Results:** Both groups showed reduced AHs after NFB, with no group differences. Compared to the Sham-NFB group, the Real-NFB group showed more reduced activation in secondary auditory cortex (AC) and more reduced connectivity between AC and cognitive control regions including dorsolateral prefrontal cortex (DLPFC) and anterior cingulate. The connectivity reduction was also observed in the Real-after-Sham-NFB condition. Secondary AC-DLPFC connectivity reduction correlated with hallucination reduction in the Real-NFB group. Replicating prior results, both groups showed reduced primary auditory cortex activation, suggesting mindfulness meditation may regulate bottom-up processes involved in hallucinations.

**Conclusions:** Our findings emphasize delivering NFB from brain regions involved in medication-resistant AHs. They provide insights into auditory cortex and cognitive control network interactions, highlighting complex processing dynamics and top-down modulation of sensory information.

## Introduction

Auditory verbal hallucinations (AHs) are a hallmark symptom of schizophrenia (SZ), experienced by approximately 60-80% of individuals with the disorder.^1,2^ AHs typically manifest as hearing voices or sounds in the absence of any external auditory stimulus and are often distressing and intrusive. The neural substrates underlying AHs in SZ involve distributed brain regions forming functional brain networks. Hyperactivity in the superior temporal gyrus (STG), the anatomical location of the auditory cortex (AC), has been implicated in the occurrence of AHs.^3–7^ According to Northoff’s resting-state hypothesis of AHs, STG hyperactivity coexists with abnormal STG interactions with the Default Mode Network (DMN) and the Central Executive Network (CEN).^8–10^ Because the DMN is normally active during rest and involved in self-referential thoughts while the CEN is associated with other cognitive functions (e.g. working memory),^11–13^ AHs may result from a complex interaction of abnormally elevated resting state activity in the STG, its hyperconnectivity with the DMN, and deficient regulation from CEN that normally enables the distinction between internally-generated and externally-induced activity.^10,14^ According to this model, a combination of hyperactivity and hyperconnectivity within and between AC and self-referential DMN in the absence of external stimuli are misidentified and perceived as AHs. Northoff’s hypothesis extends earlier hypotheses that proposed misattribution of self-generated stimuli to others instead of self,^15,16^ as well as models that suggest a breakdown in the network of brain regions involved in auditory processing and inhibitory control.^6^

AHs can be resistant, or only partially responsive, to antipsychotic medications,^17–19^ which points to the need for alternative therapeutic strategies. A promising new, non-invasive approach is real-time neurofeedback, a procedure in which individuals receive moment-by-moment feedback based on their brain signals [e.g., electroencephalography, magnetoencephalography, functional magnetic resonance imaging (fMRI)]^20^. fMRI-based real-time neurofeedback (NFB), in particular, has been promising for its ability to target specific brain regions with high spatial precision as well as to monitor multiple brain regions simultaneously, thereby providing a more comprehensive and personalized approach to brain regulation and symptom management.^21,22^ There has been encouraging progress investigating the efficacy of NFB as a treatment for AHs.^23–26^ However, most studies do not provide overt instructions or strategies to help subjects successfully regulate fMRI signals, resulting in the need for multiple training sessions for each participant.^20,27^ Clear guidance for participants with SZ is critical as this population exhibits self-regulation deficits including impaired cognitive control and emotion regulation.^28^ These impairments affect decision-making and reinforcement learning,^29–31^ making it more challenging for individuals with schizophrenia to acquire new skills. Combining mindfulness meditation and NFB, we recently developed an innovative paradigm whereby patients with treatment resistant AHs learn to modulate brain activation in the STG.^32^

Specifically, we taught participants "mental-noting" meditation to help them ignore pre-recorded sentences spoken in a stranger’s voice and downregulate STG activation. This guided neurofeedback training resulted in reduced STG activation and lower AHs scores after a single 21-minute session, suggesting this combined approach may be effective for medication-resistant AHs.^32^ Our previous study had limitations, notably the absence of a sham condition and comprehensive assessment of whole-brain impact. To address these gaps, the current sham-controlled, randomized study aims to investigate the broader brain network effects of NFB. Here we sought to examine if NFB delivered from a brain region implicated in AHs (STG) would engage cognitive control brain regions in contrast to NFB from an unrelated area.

Additionally, we aimed to replicate our previous findings showing that NFB reduces AHs and primary cortex activation.^32^ The current study has been registered with clinicaltrials.gov, updated May 17, 2024 (trial identifier: NCT05299749).

## Methods

### Overview of Study Design

Data were acquired across multiple visits: initial clinical assessment, baseline fMRI localization, two NFB visits (data related to the second NFB for the Real-NFB group are not included here), and clinical follow-up (**Figure 1A**). Participants were randomly assigned to Real- NFB or Sham-NFB groups using simple randomization for the first participant followed by alternating assignments. Initial visits included demographic collection, clinical and neurocognitive assessments, and voice recordings for subsequent fMRI sessions. Baseline fMRI comprised a T1-weighted structural scan, an STG-localizer task identifying regions sensitive to self-other voice distinction, and a resting state scan for motor cortex localization.

**Figure 1.**
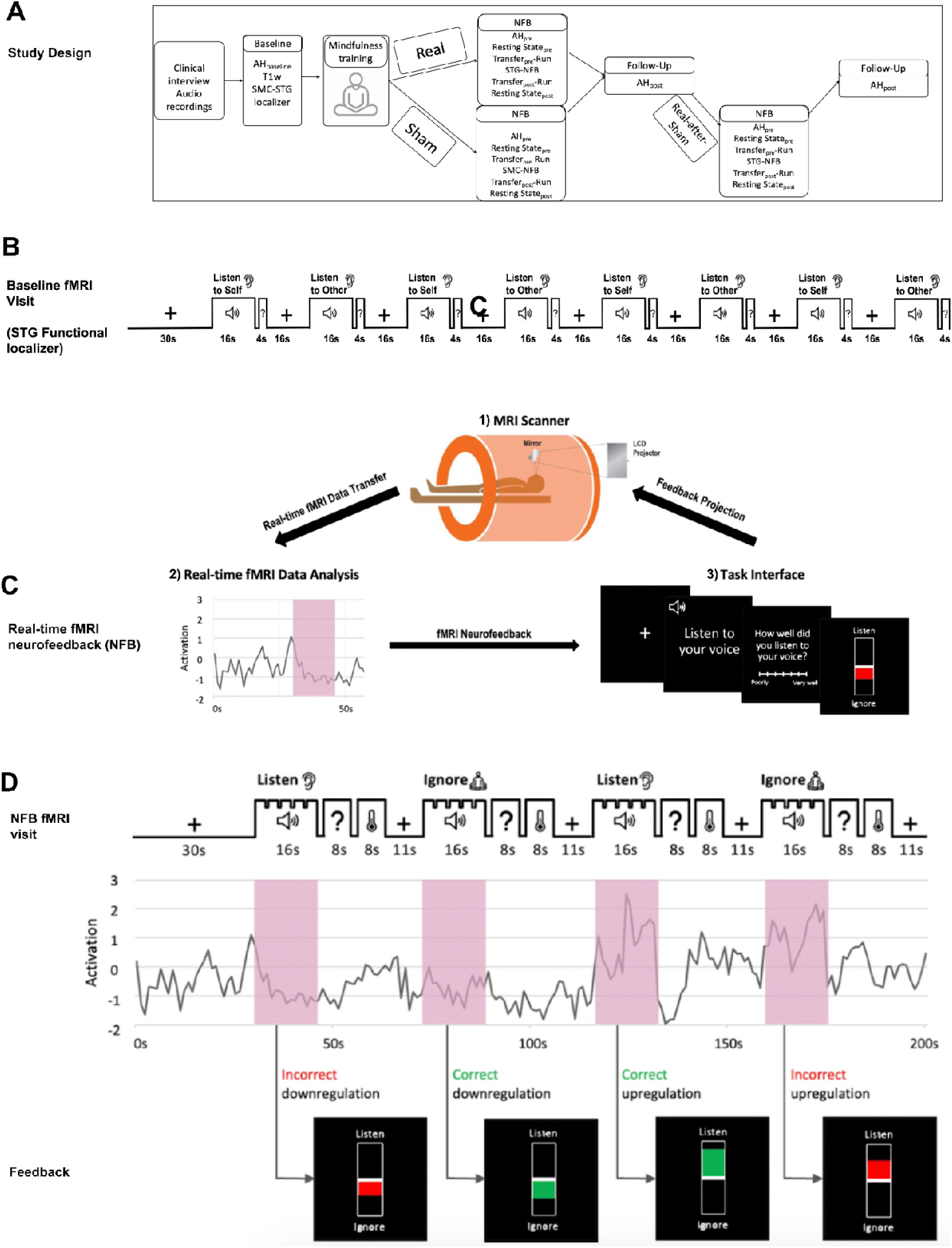
STG targeting, real-time fMRI neurofeedback experimental design (NFB). A) Overall study design. B) Baseline fMRI visit with STG functional localizer. C) Real-time fMRI neurofeedback setup that includes 1) the magnetic resonance imaging scanner, 2) real-time data analysis and 3) task Interface that forms a closed loop of information flow to the participant. D) NFB task design with an example of fMRI activation time series of the target region (superior temporal gyrus (STG) for Real-NFB and motor (MC) for Sham-NFB) which illustrates four possible types of neurofeedback that a participant could receive during the neurofeedback regulation blocks (red shadow). Green color on the thermometer indicates modulation in the correct direction (i.e., downregulation during ‘Ignore’ condition or upregulation during ‘Listen’ condition) and red color indicates modulation in the wrong direction (i.e., downregulation during ‘Listen’ condition or upregulation during ‘Ignore’ condition).

During the first neurofeedback visit, the Real-NFB group received feedback based on STG activation, while the Sham-NFB group received motor cortex feedback. The Sham-NFB group later received STG-based feedback in a second session (Real-after-Sham-NFB condition). Before and after neurofeedback, participants completed transfer tasks without feedback. Participants remained blinded to the feedback source (STG for Real-NFB and Real- after-Sham-NFB, motor cortex for Sham-NFB). All procedures were identical except for the region generating the feedback signal.

### Participants

Participants were recruited from clinical services at the Boston Veterans Affairs Healthcare System and McLean Hospital. All participants gave written informed consent obtained in accordance with the guidelines of the Institutional Review Board (IRB) of Boston VA Healthcare System and the IRB of Harvard Medical School, which served as the IRB of record for McLean Hospital, Massachusetts Institute of Technology (MIT), and Northeastern University.

Eligibility criteria included i) a diagnosis of schizophrenia or schizoaffective disorder, verified with the Structured Clinical Interview for DSM-5 (SCID-5^33^); ii) moderate or severe medication-resistant AHs at least three times a week in the past month (PANSS item #3 ≥ 4) ;iii) age 18-55 years old; iv) stable doses of antipsychotic medications; v) right-handedness (Edinburgh Handedness Inventory ≥ 60); vi) native English speakers; vii) estimated IQ ≥ 70 (WASI); and viii) normal or corrected-to-normal vision and normal hearing. Participants were excluded if they i) had a history of neurological or traumatic head injury in the past six months;ii) had alcohol use disorder or substance use disorder in the past three months (as per DSM-5) or used alcohol in the previous 24 hours; iii) were pregnant; or iv) had other MRI contraindications.

Forty-one patients enrolled in this study. After enrollment, six participants were deemed ineligible due to meeting exclusion criteria. Five participants withdrew during the course of the study. Five participants were lost to follow-up. This resulted in a total of 25 participants (36.1 ± 10.0 years; 24-54 years; 24% females) who were randomly assigned to receive either Real-NFB (n=12) or Sham-NFB (n=13). After randomization, two participants in the Sham-NFB group were lost to follow-up. Two Real-NFB participants had incomplete transfer task fMRI data after NFB. One participant’s AHs score was completed more than 200 days after the NFB session due to COVID lockdown, therefore we excluded the post-NFB AHs scores. See the CONSORT diagram (**Figure S1**) for details.

Sociodemographic and clinical characteristics are summarized in **Table 1**. A sample of 23 participants (mean age=36.34 ± 10.26; 25% females) was included for the final analysis Real-NFB (n=10) or Sham-NFB (n=13). During the course of our study, we were required to change sites and scanners. Among the 23 participants, six were scanned on a Siemens Trio scanner at MIT, 10 were scanned on a Siemens Prisma scanner at MIT and seven were scanned on a Siemens Prisma scanner at Northeastern University (for details see **Table S1**).

**Table 1.** Sociodemographic and clinical information.

|  | Real condition | Sham condition | Real vs. Sham |
| --- | --- | --- | --- |
| *N* | n=10 | n=13 |  |
| **Sociodemographic** |  |  |  |
| Age (mean/SD) | 34.6 (10.01) | 35.1 (10.65) | t-test: $p=.82$ |
| Sex: female (*n*/%) | 2/20% | 4/30.7% | Wald test: $p=.57$ |
| Race: White (*n*/%) | 6/60% | 11/84.6% | Wald test: $p=.19$ |
| Education [years] (mean/SD) | 13.7 (3.4) | 14.0 (2.0) | t-test: $p=.63$ |
| IQ: WASI (mean/SD) | 99.5 (21.2) | 97.3 (16.9) | t-test: $p=.44$ |
| **Auditory Verbal Hallucination Severity (mean/SD)** |  |  |  |
| PSYRATS |  |  |  |
| Baseline | 25.16 (6.99) | 26.25 (7.08) | t-test: $p=.82$ |
| Post-NFB | 21.66 (9.0) | 23.5 (6.56) | t-test: $p=.45$ |
| Post-NFB* | 19.5 (5.0) | 22.62 (6.75) | t-test: $p=.52$ |

*Note.* All Real vs. Sham tests were two-tailed independent samples tests. PSYRATS=The Psychotic Symptom Rating Scales. WASI=Wechsler Abbreviated Scale Intelligence (Full Scale) NFB=real-time fMRI neurofeedback, \*NFB=Real-after-Sham-NFB.

### Clinical Assessments

We administered the SCID-5 to verify diagnoses of schizophrenia or schizoaffective disorder. To measure the severity of AHs, which was our primary symptom of interest, we administered the auditory hallucinations subscale of the Psychotic Symptom Rating Scale (PSYRATS-AHs^34^). We administered the PSYRATS-AH at the baseline clinical visit, at the beginning of every fMRI study visit, and also at the clinical follow-up visit. All baseline clinical and neuropsychological evaluations were conducted in-person.

### Audio recording and processing

Patients recorded 170 sentences to be used in the ‘self-voice’ condition on an Olympus, digital recorder, model VN-722PC. Recording was carried out in a sound-proof room and sentences were read without emotional intonation. All sentences had a structure of subject + verb + adjective + object (e.g., “Jane liked chocolate chip cookies”) with an average length of 6.65 words (SD=1.13). All sentences were neutral in affect and written in third person. Recorded sentences were processed in 3 steps: 1. Audio intensity was normalized using Praat (http://www.fon.hum.uva.nl/praat/). 2. Background noise was removed using Audacity (http://www.audacityteam.org). 3. Individual sentences were segmented using Praat. The same sentences were recorded by a male in his 40s and edited in the same manner to be used in the ‘male other’ condition. We further processed the ‘male other’ sentence in two ways to mimic female voice: 1) We changed the formant shift ratio to 1.2 to account for a shorter vocal tract. 2 We changed the pitch median to 220 Hz to account for greater vocal fold tension.

### Baseline STG Functional Localizer

Participant-specific STG was defined during the baseline visit using a functional localizer task (Self-Other Task; **Figure 1B**) in which participants listened to pre-recorded sentences.

Eighty unique sentences were presented: 40 sentences in the subject’s own voice and 40 in a sex-matched stranger’s voice. The Self-Other Task included four blocks each of 16 s duration (i.e., two for self-voice, two for other-voice) interspersed with fixation blocks of the same length. Each ‘self-voice’ block included five unique sentences spoken in the subject’s voice; each ‘other-voice’ block included five unique sentences spoken in a sex-matched stranger’s voice. For each subject, sentences in self-voice and other-voice did not overlap. The fixation blocks included a silent block where participants gazed at a crosshair in the center of the screen. To help sustain attention, after each voiced block, subjects were prompted with a ‘yes’ or ‘no’ question on the screen regarding the content of the last sentence they had heard in the block (e.g., “Did Jane like chocolate chip cookies?”) and submitted their response via button press. After the fixation block, they answered a real-world question (e.g., “Is the White House in Washington D.C.?”). These blocks were split evenly across two functional runs (4 minutes and 26 seconds each). Each run contained 12 pseudo randomized blocks (4 blocks per condition) such that fixation blocks did not appear back to back. In an off-line analysis (see fMRI Data Preprocessing), a participant’s specific target region of interest (ROI) was defined for the thresholded contrast self-voice>other-voice in bilateral STG (MNI space). From the resulting cluster the top 100 voxels were used as the final ROI.

### Baseline MC Functional Localizer

We localized each participant’s MC using resting state fMRI (rs-fMRI). MRI acquisition details can be found in supplementary materials. Preprocessing of rs-fMRI data was performed in FSL 6.0^35^ and included: motion correction, brain extraction, co-registration, smoothing and bandpass filtering (see more details^36^). We performed an independent components analysis (ICA) on the preprocessed functional scans using Melodic ICA version 3.14^37^ with dimensionality estimation using the Laplace approximation to the Bayesian evidence of the model. We compared each of the ∼35 spatiotemporal components to the spatial map of the MC derived from rs-fMRI of ∼1000 participants^38^ using FSL’s “fslcc” tool and selected the ICA component that yielded the highest spatial correlation for each participant. These ICA components were thresholded to select the upper 10% of voxel loadings and then binarized to obtain participant-specific MC masks. Visual inspection was performed and all components maps were determined to be satisfactory in covering canonical MC brain regions.^39^ For details on MRI Data Acquisition see supplemental material and **Table S1**.

### Mental-noting training

We trained participants on a mindfulness technique called “mental-noting” (from Vipassana insight meditation^40^) which consists of two main components: “concentration” and “observing sensory experience”. The experimenter first explained to the participants that mental- noting entails being aware of one’s sensory experience without engaging in or dwelling on the details of the content; in other words, one would “note” the sensory experience (e.g., “hearing”, “seeing”, “feeling”) at the forefront of their awareness and then let it go after it has been noted. The experimenter also introduced the concept of an “anchor”, or a sensory experience to which one could easily switch their attention, such as breathing or sensations on the left toe.

Participants were encouraged to use their personal anchors when they noted consecutive “hearing”. The experimenter demonstrated noting out loud by verbalizing the predominant sensory modality approximately once per second. Participants were then asked to practice mental-noting out loud to demonstrate the ability to describe sensory awareness without engaging in the content and stop consecutive “hearing”. We instructed participants to use the mental-noting strategy during the “other-voice” blocks in order to help them ignore the other voice and all sounds. Participants were trained outside the scanner just before the NFB scan. The training lasted about 15 min and all participants were able to grasp the concept within that time. The training was done shortly after the baseline PSYRATS assessment and shortly before entering the MRI scanner (see supplement for mindfulness-training script).

### Real-NFB task

NFB was delivered in the form of a thermometer with the color green representing positive-feedback and the color red representing negative-feedback. The extent of the thermometer represented brain activation. NFB setup includes three main components that form a loop of information flow (**Figure 1C**). All participants randomized to Real-NFB (n=10) heard a set of 80 unique sentences (average length=6.64 words, SD=1.31) (a different set of sentences than the ones presented during the localizer scan) presented via earphones in the scanner.

Forty sentences were in their own voice and 40 in a stranger’s voice. These sentences were not significantly different in their length from the 80 sentences heard in the STG localizer Self-Other Task [*t*(158)=0.34, p=0.73]. There were six runs for this task, each lasting 2.5 minutes. The first and last runs were transfer runs (i.e., they did not provide NFB; NFB-transfer-pre and -post), while runs 2–5 provided real-time NFB from individualized STG masks. Each run had four randomized blocks composed of two listen blocks and two ignore blocks, each lasting 16 s (**Figure 1D**). Before each block, participants saw a prompt stating either “listen to self” to signal that they were asked to attend to the sentences played over the earphones (resulting in up- regulating STG) or “ignore all sounds”, to signal that they were asked to ignore the sentences along with any other environmental noises including the scanner noise (resulting in down- regulating STG). To successfully ignore the other-voice sentences, participants were instructed to use the mental-noting strategy. After each feedback block, participants saw a ‘thermometer’ showing their activation level in the STG. The height of the ‘thermometer’ bar reflected a median activity level for a given block. In addition, to gauge the participant’s assessment of their own performance during the tasks, after each listen block, a prompt asked the participant to rate how well they were able to attend to the voices on a scale of 1 (completely attended to) to 6 (could not attend at all); after each ignore block, a similar prompt asked how well they were able to ignore all sounds. Participants responded to these prompts via button press. The STG estimates were calculated as the median activity across all voxels within each participant’s STG mask (as defined by STG Functional Localizer) and co-registered to the current fMRI volumes. To accomplish the voxel-wise estimation in real-time,^41^ we first collected 30 seconds of baseline data and then continuously performed an incremental general linear model (GLM) fit with subsequent incoming images. This method accounts for the mean voxel signal and linear trends. To discount components of the voxel signal due to nuisance sources (e.g., low-frequency signal drifts), the GLM reconstruction of the expected voxel intensity at time t was subtracted from the measured voxel intensity at time t, leaving a residual signal that has components due to two sources: BOLD signal fluctuations and unmodeled fMRI noise. This residual was scaled by an estimate of voxel reliability, which was computed as the average GLM residual over the first 25 functional images of the baseline. This analysis resulted in an estimate of the strength of activation at each voxel at time t in units of standard deviation.

### Sham-NFB task

All participants randomized to Sham-NFB (n=13) completed the feedback task that was structured identically to the Real-NFB with the exception that feedback was provided from the personalized MC. All other instructions and parameters were the same.

### fMRI Data Preprocessing for mask generation and analysis

Preprocessing and data analyses were completed using FSL Version 6.0.3.^35^ Preprocessing included brain extraction using BET (Brain Extraction Tool), motion correction using MCFLIRT (intramodal motion correction tool.^42^ The remaining fMRI signals were spatially blurred with 6-mm full-width-at-half-maximum (FWHM) Gaussian kernel. A subject-dependent number of individual nuisance regressors for removing outlier time points were created using the fsl_motion_outliers tool, employing the DVARS metric. This metric applies a threshold on the root-mean-squared intensity difference of volume N to volume N+1.^43^ The threshold used was the standard boxplot outlier threshold set in FSL.^43^ Participants were excluded if they had >20% of movement outliers.^44^

### Clinical Analysis, effect of NFB on PSYRATS-AH scores

Statistical tests for PSYRATS-AH data were conducted using R Studio Version 1.0.136.^45^. Statistical significance level was set at .05.

### Brain Activation Analyses

Activation analyses were restricted to the bilateral anterior and posterior temporal lobes as defined by the Harvard-Oxford atlas^46^ with a threshold at 85% probability to remove any non temporal lobe voxels as an *a priori* ROI based on a previous report.^32^ We examined how temporal lobe activation changes after NFB (pre-post) differed between Real-NFB and Sham- NFB while participants were using the nothing practice to ignore the stranger’s voice (other-voice>fixation-block, hereafter ‘ignore-other-voice’).

### Functional Connectivity Analyses

To examine functional connectivity in Real-NFB relative to Sham-NFB, we performed a generalized psychophysiological interaction (gPPI) analysis.^47,48^ Specifically, we examined the functional connectivity of a secondary auditory cortex (AC) cluster identified from the activation analysis during the *ignore-voice* blocks (depicted in Blue in **Figure 3A**, peak: *X*=56, *Y*=-24, *Z*=- 6, BA 22). First-level GLM analyses included four regressors: psychological, physiological, gPPI, and nuisance. *Listen-voice-* and *ignore-voice*-*block* durations comprised the psychological regressor, modeled as a boxcar function, convolved with a single gamma hemodynamic response function. For each participant, we extracted the physiological regressor as the time series for the functional cluster identified in the two-way mixed effect ANOVA during the *ignore- voice-block* contrast across groups. The gPPI regressor was the product of the demeaned physiological regressor and the psychological regressor, which was zero-centered between the minimum and maximum values. The following nuisance regressors were modeled: global mean time series of each preprocessed run, six motion parameters, and motion outliers. These individual contrast images were then entered into a second level fixed-effects analysis including both runs for each participant. Mixed-effect analysis of variance tests with factors of Time (pre-NFB vs. post-NFB) x Group (Real-NFB vs. Sham-NFB) on each participant’s STG and whole brain connectivity Z scores were conducted. Significant main effects and interactions were followed up with post hoc testing.

### Statistical Tests

Results are reported at p-FWE<0.05 with small volume correction for bilateral temporal lobes (Harvard-Oxford atlas).^46,49,50^ Whole-brain and gPPI analyses used FSL randomize^51,52^ with 5,000 permutations, automatic outlier deweighting, and probability threshold-free cluster enhancement.^53^ Results are thresholded at z>3.1 (p<0.001).

### Replication Analyses

We sought to replicate our previous finding^32^ of reduced primary auditory cortex activation (MNI:60,-18,10; BA41) using a 10mm sphere ROI. Self-voice vs. other-voice contrasts were calculated per run, with scanner as a nuisance regressor. We correlated hallucination scores with ROI activation during ignore-voice blocks. Details in supplement.

## Results

### Changes in PSYRATS-AH

Paired t-tests within the groups showed improvements in AHs, as measured by the PSYRATS-AH, from pre- to post-intervention for both Real-NFB [*t*(10)=1.87, *p*=.04, Cohen’s *d*=.82)] and Sham-NFB [*t*(10)=1.90, *p*=.04, Cohen’s *d*=.48]. Using a two-way ANOVA to investigate the effects of Real-NFB vs. Sham-NFB on PSYRATS-AH, we found a significant main effect of Time [F(2, 18)=2.62, *p*=.01, Cohen’s *f^2^*=.42], but no effect of Group [F(2, 18)=1.33, *p*=.19] or Time x Group [F(2, 18)=1.62, *p*=.12] interaction (**Figure 2A**).

**Figure 2.**
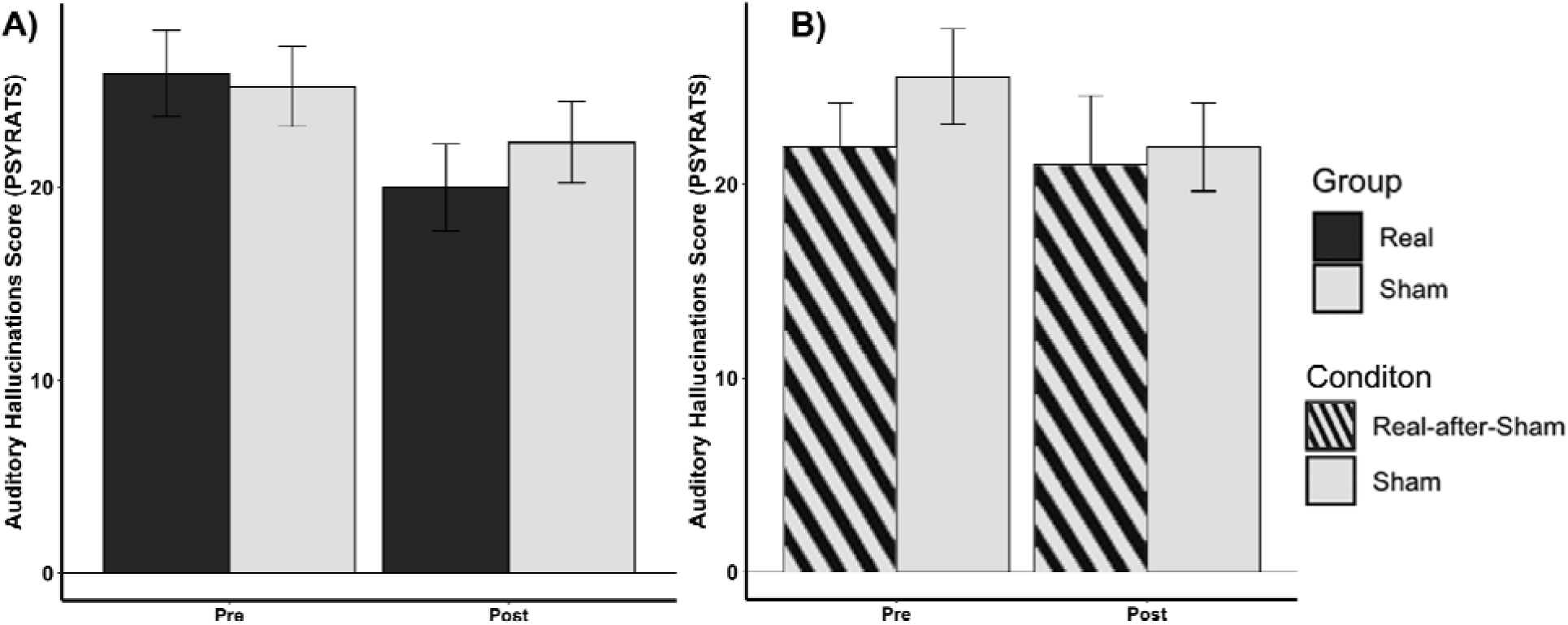
Auditory hallucinations score changes post-NFB. (A) AH-Scores pre- and post- NFB for Real-NFB and Sham-NFB groups. (B) AH-Scores pre- and post-NFB for Real-after- Sham-NFB and Sham-NFB conditions. Auditory hallucinations were assessed using the Psychotic Symptom Rating Scale (PSYRATS^54^). All bars reflect mean and all error lines reflect standard error of the mean.

**Figure 3.**
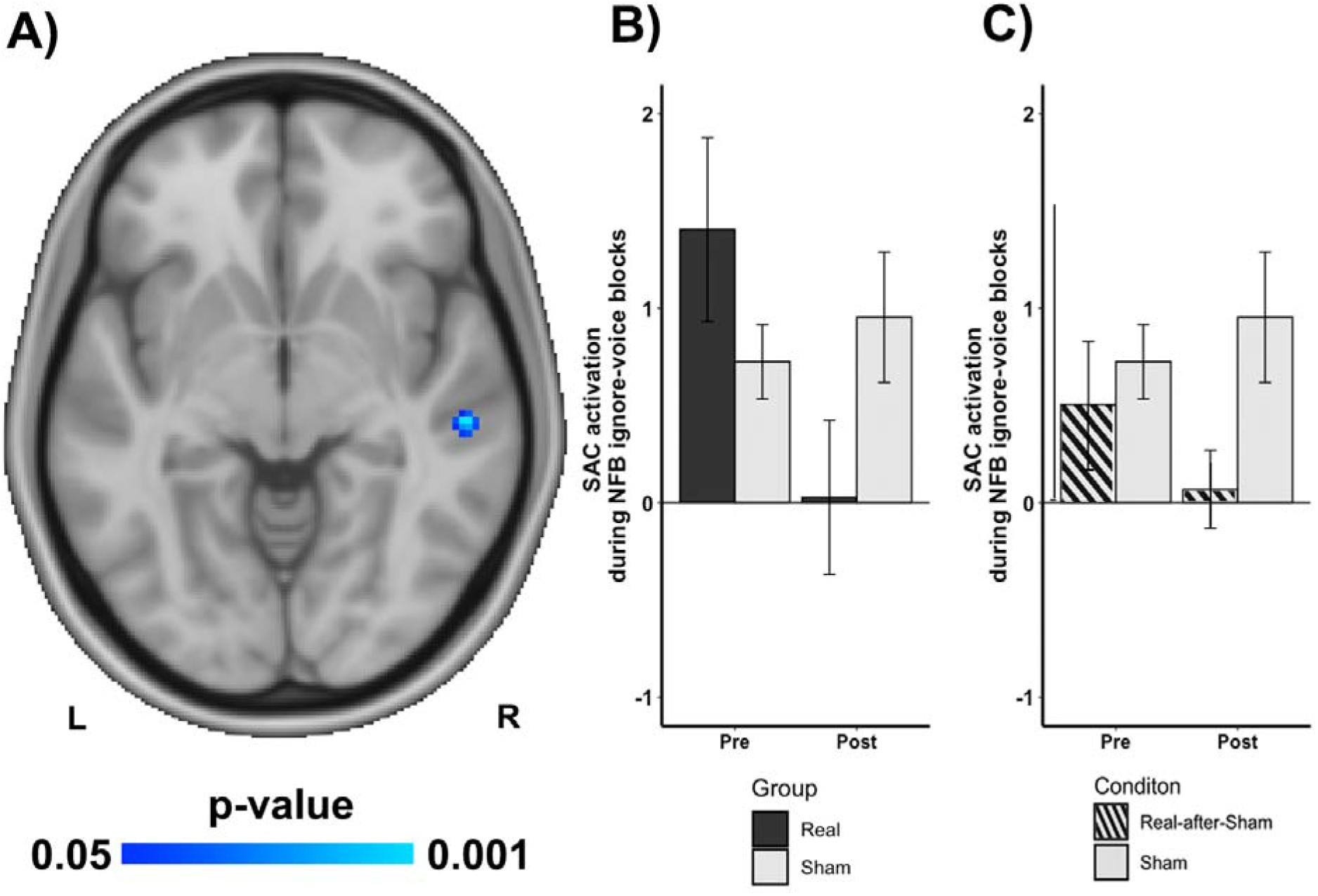
Secondary auditory cortex activation changes post-NFB. (A) Region exhibiting significant Time x Group differences in secondary auditory cortex (BA22) activation during *ignore-voice* task and Real or Sham mindfulness enhanced neurofeedback (NFB). (B) Significantly decreased activation pre-to-post intervention in the Real-NFB group relative to the Sham-NFB group. (C) Significantly decreased activation pre-to-post intervention in the Real- after-Sham-NFB relative to the Sham-NFB condition. All bars reflect mean and all error lines reflect standard error of the mean; statistics are nonparametric and FWE-pTFCE small volume corrected. Secondary auditory cortex (SAC).

Similarly, a repeated measures ANOVA comparing Real-after-Sham-NFB and Sham- NFB, showed a significant main effect of Time [*F*(2, 16)=2.61, *p*=.01, Cohen’s *f^2^*=1.14], with no effect of Condition [F(2, 16)=.92, *p*=.37] or Time X Condition [F(2, 16)=1.05, *p*=.30] interaction (**Figure 2B**). No additional reductions in AHs were observed in Real-after-Sham-NFB [*t*(8)=.96, *p*=.18, Cohen’s *d*=.10]. See mean and standard deviation values for pre- and post-intervention in **Table S2**.

### Neuroimaging Analyses Motion

We did not exclude any participants due to motion as no participants exceeded 20% of high motion outliers.^55^ We did not find any difference when contrasting motion between Real- NFB and Sham-NFB groups [*F*(2, 18), *p*>3, Cohen’s *f^2^*<.1] or between Real-after-Sham-NFB and Sham-NFB conditions [*F*(2, 19)<.90, *p*>3, Cohen’s *f^2^*<.1].

### Effect of NFB on brain activation during ignore-voice

Using a two-way ANOVA, we tested whether brain activation in the voxels of the left and right temporal poles changed while participants were using the mental-noting practice to ignore the strangers voice. During the *ignore-voice* blocks, there was a significant Time (pre- and post-intervention) X Group (Real-NFB and Sham-NFB) interaction in the secondary AC (*n*=23; *X*=56, *Y*=-24, *Z*=-6, BA22; non-parametrically SVC at *p*<.001, Cohen’s *f^2^*=.54; **Figure 3A**), such that there was a greater reduction in right secondary AC activation after Real-NFB vs. after Sham- NFB. No main effects were found for Group or Time (*p*>.1). Post hoc two-sample t-tests between the groups showed no significant difference between groups at pre-NFB for right secondary AC activation during *ignore-voice* (*t*(18)=1.52, *p*=.15, Cohen’s *d*=.67, **Figure 3B**).

Paired t-tests within the groups only showed lower secondary AC activation for the Real-NFB group during the *ignore-voice* task in the secondary AC at post-NFB (*t*(9)=2.7, *p*=.01, Cohen’s *d*=1.05, **Figure 3B**) with no significant difference for the Sham-NFB (*t*(11)=.78, *p*=.77, Cohen’s *d*=.23, **Figure 3B**) or Real-after-Sham (*t*(11)=1.1, *p*= .13, Cohen’s *d*=.45, **Figure 3C**) conditions. However, there was a trending Condition effect between Real-after-Sham-NFB and Sham-NFB (*F*(2, 19)=1.83, *p*=.08, Cohen’s *f^2^*=.41; **Figure 3C**), such that right secondary AC activation was further reduced in the Real-after-Sham condition following Sham-NFB. No main effects were found for Time or Condition [Fs(2, 19)<.90, *ps*>3)]. This was confirmed by post hoc paired t-tests between Real-after-Sham-NFB and Sham-NFB conditions showing no significant difference between conditions at pre-NFB for secondary AC activation for *ignore-voice* [t(10)=.52, *p*=.60, Cohen’s *d*=.07] but a significantly lower activation for *ignore-voice* in the Real- after-Sham-NFB than the Sham-NFB condition at Post-intervention [t(16)=2.2, *p*= .05, Cohen’s *d*=.74]. See mean and standard deviation values for pre- and post-intervention in **Table S2**.

### Effect of NFB on functional connectivity during ignore-voice

When we used gPPI analysis to examine if NFB produced changes in functional connectivity between secondary AC and the rest of the brain, we found a significant Time X Group interaction of change in functional connectivity between secondary AC and right dorsolateral prefrontal cortex (rDLPFC, *n*=22: *X*=28, *Y*=50, *Z*=22; BA10; non-parametrically at *p*<.001, Cohen’s *f^2^*>1.56; **Figure 4A**). Functional connectivity between secondary AC and rDLPFC during *ignore-voice* reduced for the Real-NFB group while it increased for the Sham- NFB group. This was confirmed by post hoc two-sample t-tests between the groups showing no significant difference at pre-NFB for secondary AC functional connectivity to the rDLPFC during *ignore-voice* (*p*>.3,Cohen’s *d*<.1, **Figure 4B**), but significantly lower secondary AC functional connectivity to the rDLPFC for *ignore-voice* in the Real-NFB group than the Sham-NFB group at post-NFB [*t*(18)=4.42, *p*=.00005, Cohen’s *d*=1.89, **Figure 4B**). Additionally, there was a significant interaction effect for Condition (*F*(2, 19)=2.25, *p*=.056, Cohen’s *f^2^*=1.63) of change in functional connectivity between secondary AC and rDLPFC (**Figure 4C**). The pattern of this interaction suggests that functional connectivity between secondary AC and rDLPFC during *ignore-voice* remained similarly reduced after receiving Real-after-Sham-NFB, while showing a trend towards increased connectivity in the Sham-NFB condition. This was confirmed by post hoc two-sample t-tests between the conditions showing a trending difference at pre-NFB for secondary AC functional connectivity to rDLPFC during the *ignore-voice* block (*t*(16)=1.72, *p*=.09, Cohen’s *d*=.67, **Figure 4C**), but a significantly lower functional connectivity for *ignore- voice* in the Real-NFB condition than in the Sham-NFB condition at Post-intervention(*t*(16)=2.74, *p*=.01, Cohen’s *d*=1.07, **Figure 4C**). See mean and standard deviation values for pre- and post-intervention in **Table S2**.

**Figure 4.**
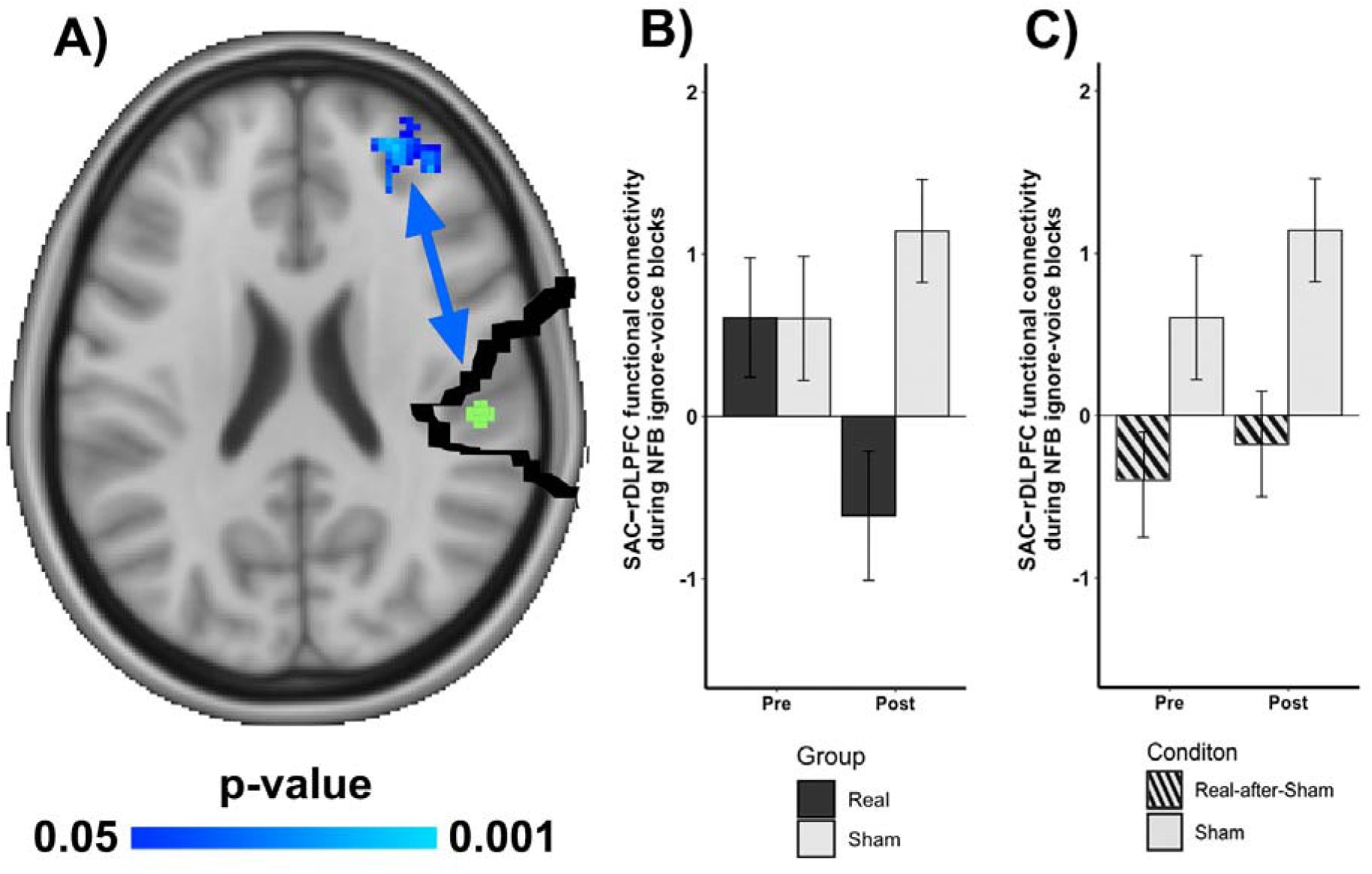
Whole brain analysis for functional connectivity changes of secondary auditory cortex post-NFB. (A) Regions exhibiting significant Time x Group differences in secondary auditory cortex (SAC: green ROI seed in the superior temporal gyrus, peak: 56, -24, 6; BA22) functional connectivity and right dorsolateral prefrontal cortex during *ignore-voice* condition and Real or Sham mindfulness enhanced neurofeedback condition (NFB). (B) Significantly decreased SAC-rDLPFC functional connectivity pre-to-post intervention in the Real-NFB group relative to the Sham-NFB group. (C) Significantly decreased activation pre-to-post intervention in the Real-after-Sham-NFB condition relative to the Sham-NFB condition. All bars reflect mean and all error lines reflect standard error of the mean; statistics are nonparametric and FWE- pTFCE small volume corrected. rDLPFC=right dorsolateral prefrontal cortex; SAC=Secondary auditory cortex.

### Effect of NFB on activation in cognitive control regions during ignore-voice

When assessing the activation change for the rDLPFC obtained during gPPI analyses during the *ignore-voice* blocks, there was a significant Time X Group interaction [*F*(19)=2.49, *p*=.02, Cohen’s *f^2^*=.93, **Figure 5A**] such that activation for the rDLPFC during *ignore-voice* was maintained in the Real-NFB group while activity significantly decreased in the Sham-NFB group. This was confirmed by post hoc paired t-tests within the conditions showing no significant difference from pre- to post-intervention for *ignore-voice* in the Real-NFB group [*t*(9)=1.15, *p*=.13, Cohen’s *d*=.22, **Figure 5B**] while significantly decreased in the Sham-NFB group at post-intervention [t(12)=1.19, *p*=.003, Cohen’s *d*=1.30 **Figure 5B**].

**Figure 5.**
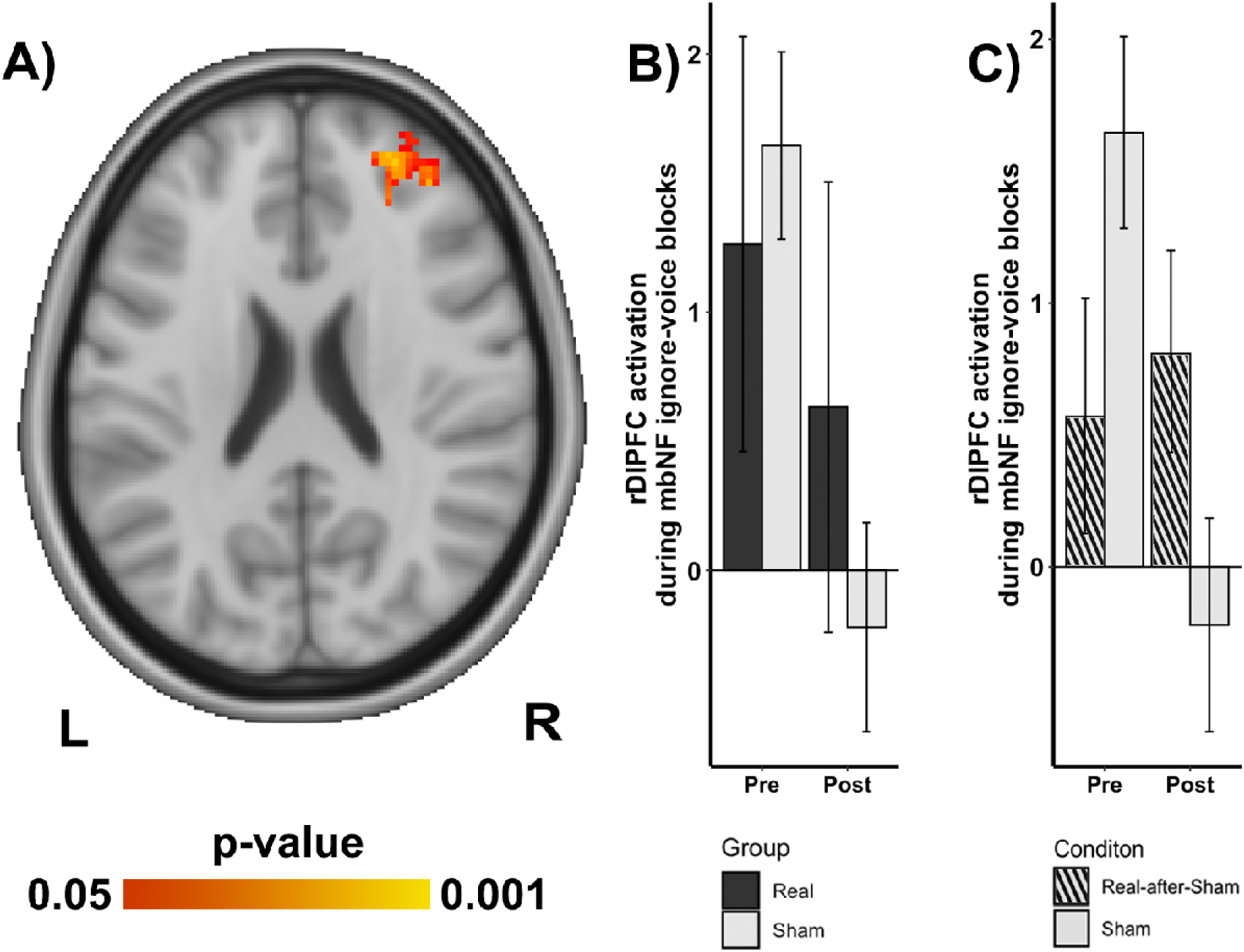
Activation changes in cognitive control brain regions post-NFB. (A) Region in the right dorsolateral prefrontal cortex (rDLPFC) where activation reflects a significant Time x Group interaction. (B) rDLPFC maintained similar activation pre-to-post intervention in the Real- NFB group relative to a significant decrease in the Sham-NFB group. (C) rDPFC maintained similar activation pre-to-post intervention in the Real-after-Sham-NFB condition relative to a significant decrease in the Sham-NFB condition. All bars reflect mean and all error lines reflect standard error of the mean; statistics are nonparametric and FWE-pTFCE small volume corrected.

Similarly, although there was no Time X Condition interaction for the rDLPFC (F(19)=.71, *p*=.48, Cohen’s *f^2^*=.21; **Figure 5C**), the activation for the rDLPFC during *ignore-voice* was maintained after receiving Real-after-Sham NFB while activity significantly decreased in the Sham-NFB condition. This was confirmed by post hoc paired t-tests within the conditions showing no significant difference from pre- to post-intervention for *ignore-voice* in the Real-after- Sham-NFB condition (t(10)=.38, *p*=.64, Cohen’s *d*=.16, **Figure 5C**). See mean and standard deviation values for pre- and post-intervention in **Table S2**.

### Relationship between PSYRATS-AH changes and changes in activation and functional connectivity during ignore-voice

When assessing the relation between the changes in activation of the secondary AC and the change in AHs score from pre- to post-intervention, we found no relationship for any group or condition. When assessing change in functional connectivity between secondary AC and right-DLPFC and change in AHs score from pre- to post-intervention, we found a trending relationship between reduction in secondary AC-rDLPFC connectivity and reduction in AHs scores in the Real-NFB group (n=10, r=.45, *p*=.09; **Figure 6**), while there was no relationship for the Real-after-Sham-NFB condition (n=11, r=.005, *p*=.49; **Figure 6**) and a negative correlation for Sham-NFB group (n=12, r=-.24, *p*=0.78). A post hoc Fisher’s z test to compare between these correlations revealed a trend at *p*=.06.

**Figure 6.**
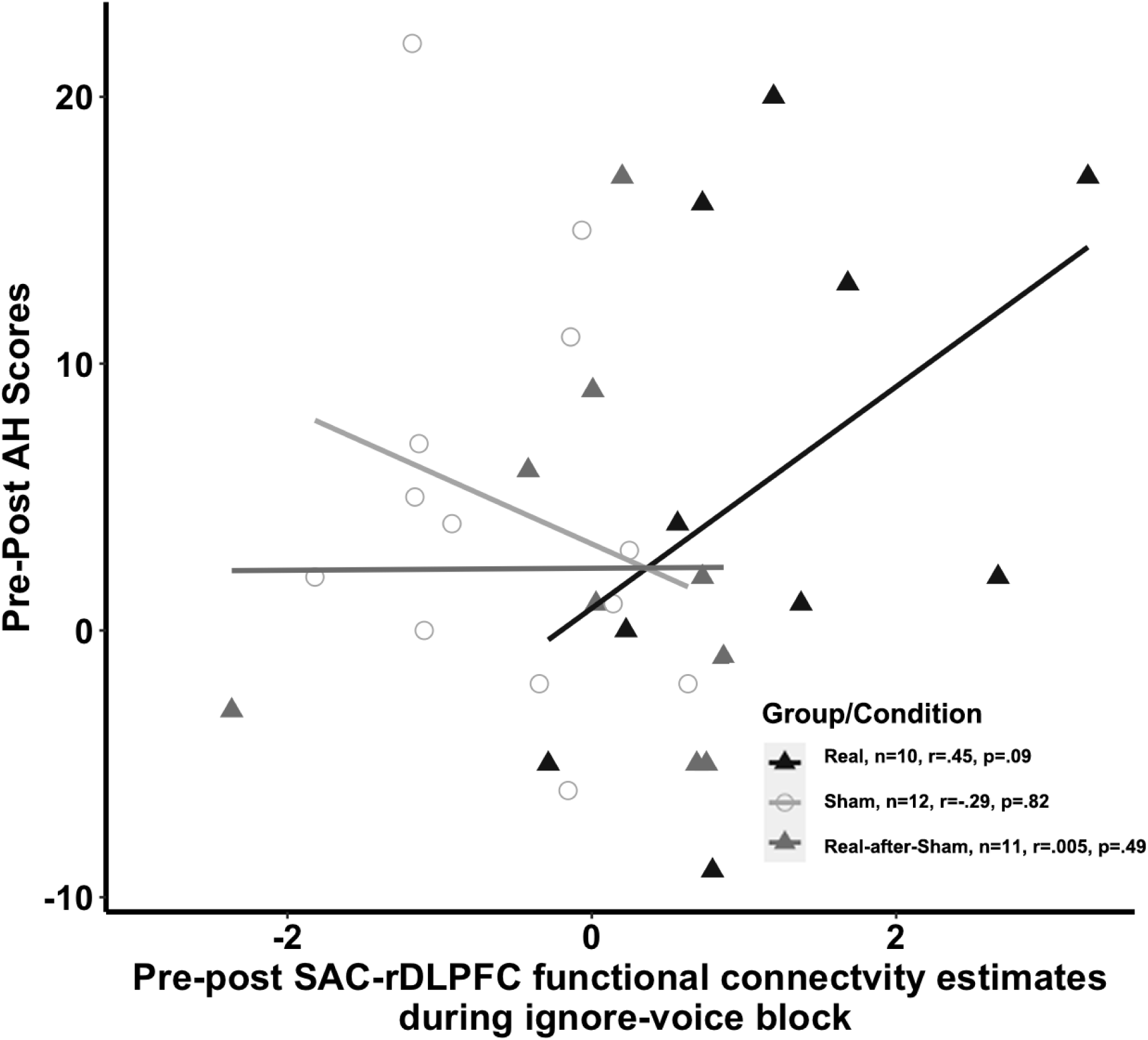
Changes in AH-scores and secondary auditory cortex functional connectivity post-NFB. Correlations between pre-post change in Auditory Hallucinations Scores (AHs) and pre-to-post change in secondary auditory cortex (SAC) and right dorsolateral prefrontal cortex (rDLPFC) functional connectivity for Real-NFB (black triangles), Sham-NFB (open circles) and Real-after-Sham-NFB (gray triangles).

## Discussion

In this study, we examined if real-time-fMRI-neurofeedback (NFB) delivered from a brain region implicated in auditory verbal hallucinations (AHs), i.e., superior temporal gyrus (STG; Real-NFB), combined with “mental-noting” practice (a type of meditation), would lead to changes in activation, functional connectivity, and AHs scores relative to a Sham-NFB delivered from an unrelated brain area, i.e., motor cortex (MC). We found that only the Real-NFB condition showed a significant reduction in 1) activation of the secondary auditory cortex (AC), 2) activation of a cognitive control region (i.e., rDLPFC), and 3) functional connectivity between secondary AC and rDLPFC. AHs were reduced independent of group assignment. We also found a trend-level relationship between the reduction in secondary AC-rDLPFC connectivity and reduction in AHs-scores, in the Real-NFB group only. Finally, in a set of ROI-based replication analyses (see supplemental material), we replicated that NFB reduced primary AC activation, which is in line with our previous finding.^32^

Both Real-NFB and Sham-NFB groups showed reduced auditory hallucination scores after the intervention. Since both groups practiced mental-noting during neurofeedback training (discussed below), these reductions may be attributed to meditation-related mechanisms such as increased self-awareness or expectancy effects.^56,57^ For a detailed analysis of the relationship between primary auditory cortex and auditory hallucinations, please see our replication analysis in the supplement.

The reduction in secondary AC activation in Real-NFB represents a significant finding for understanding and treating AHs. Secondary AC (BA22), a component of Wernicke’s area,^58,59^ is crucial for auditory processing,^58^ word-form recognition and language comprehension.^60,61^ Hyperactivation of secondary AC has been reported in individuals experiencing AHs.^62,63^ While secondary AC is not typically considered a part of the DMN, functional interactions between auditory regions and DMN are linked to internally-directed cognition and autobiographical memory retrieval.^64–66^ DMN contributes to self-referential processes like autobiographical memory and social cognition through interactions with medial prefrontal and posterior cingulate cortices.^65,66^ Dysfunction in DMN and increased secondary AC activity underlies impairments in self-referential thinking in SZ patients^67,68^ and contributes to AHs.^9,66^ Northoff’s resting state hypothesis proposes elevated auditory cortex activity and reduced top-down modulation as mechanisms for AHs.^9^ This hypothesis suggests a failure to distinguish intrinsic secondary AC activity from external activity. The combination of elevated intrinsic activity in secondary AC and DMN, coding inner speech, and impaired DLPFC control result in misattributing internal activity to external sources. NFB may effectively modulate activity in this region, disrupting AHs’ neural processes. This aligns with studies showing reduced auditory activation following successful AHs treatments.^32,36,69^

Our finding of a reduction in secondary AC activation was coupled with sustained activation of the DLPFC in the Real-NFB condition in contrast to DLPFC deactivation in the Sham-NFB condition. The latter may represent a response to inconsistent feedback, wherein the observed feedback is not in concordance with the perceived effort expended during Sham- NFB.^60^ Conversely, during feedback that is congruent with perceived effort during Real-NFB, sustained activation in the DLPFC may regulate the hyperactivity in auditory cortices,^6,61^ thereby restoring normal self-monitoring processes^64,70^ and reducing AHs.^62,63^

Reduced functional connectivity between the secondary AC and rDLPFC observed in the Real-NFB condition has further implications for understanding and treating AHs. It has been proposed that abnormally increased connectivity between speech-related network (including secondary AC)^58,59^ and executive network (including DLPFC) could reflect increased attentional resources for salient internal events (e.g., AHs) and inefficient top–down suppression.^65^ Prior studies have provided evidence of increased connectivity between the secondary AC and the central executive network (CEN) in individuals experiencing AHs.^62,62,66^ Our finding of decreased connectivity between secondary AC and DLPFC after Real-NFB further supports the role of brain regions involved in auditory processing and executive control play in improving AHs.^12,24,36^ This change in functional connectivity could influence AHs through several potential mechanisms. Previous research, such as that by Wolf et al.,^65^ has demonstrated that hyperconnectivity between frontal and temporal cortices in individuals experiencing AHs may reflect increased attentional resources for salient internal events, coupled with inefficient top-down suppression and executive control.^23,24,36^ Reduced STG-rDLPFC connectivity may disrupt the abnormal neural patterns underlying AHs, potentially by diminishing the attentional resources allocated to internal auditory events.^24,32,36^ However, it is important to note that without a healthy control group, we cannot definitively establish what constitutes a "normal" level of connectivity between secondary AC and DLFPC.^36,71,72^

Moreover, the trend-level relationship between reduced secondary AC-rDLPFC connectivity and decreased AHs scores in the Real-NFB condition, while not statistically significant, is noteworthy. It suggests that the neurofeedback intervention may specifically target neural circuits relevant to AHs^24,67,73^ and identifies secondary AC-rDLPFC connectivity as a biomarker for treatment response.^23^ This trend level effect provides preliminary evidence linking the changes in brain connectivity to clinical improvement,^24,36^ thus highlighting the individual variability in neurofeedback response and the need for personalized treatment approaches.^27,74^ These findings tentatively align with models of AHs emphasizing disrupted connectivity between auditory processing and cognitive control regions,^24,62,67^ suggesting that neurofeedback may potentially reconfigure these functional connections (though our interpretation remains limited by the absence of a healthy control group). However, future research with larger sample sizes and long-term symptom tracking is needed to confirm and extend the significance of these findings.

Mental-noting was practiced across Real-NFB and Sham-NFB groups. The reduction in AHs and PAC activation across participants may be attributed to this shared approach, aligning with research showing that mindfulness can modulate sensory processing.^75–78^ However, non-specific mechanisms like placebo effect or subject expectation may also play a role.^79^ Mental- noting involves labeling present-moment experiences without elaboration, thereby cultivating awareness and non-judgmental observation.^80^ These aspects may be relevant, as they potentially reduce emotional reactivity and distress associated with AHs.^81,82^ The selective engagement of secondary AC and rDLPFC in the Real-NFB group suggests interplay between neurofeedback and mindfulness techniques. While non-specific factors may also influence primary sensory areas, we suspect noting-practice contributed to its modulation^75,76,83^ irrespective of neurofeedback type; however this needs systematic exploration. Moreover, mental-noting with Real-NFB appears to facilitate specific modulation of secondary AC.^84,85^ Present-moment focus might enhance participants’ engagement with the neurofeedback task, improving attention and reducing mind-wandering.^76,86–88^ This combination may represent a novel approach to addressing neural dynamics underlying AHs, providing "top-down" regulation alongside specific neuromodulation.^89^ This approach of leveraging mindfulness benefits with targeted neuromodulation may offer a more comprehensive treatment strategy for AHs than either alone.

While we focused on therapeutic brain changes related to Real-NFB, the effects of Sham-NFB also should be carefully considered. When feedback signals are incongruent with brain states, these misaligned cues may actively interfere with treatment, creating a nocebo effect.^90^ This interference aligns with principles of neurofeedback training, which relies on operant conditioning where brain activation patterns are reinforced through contingent feedback signals.^82,91^ Studies have shown incongruent feedback can impair learning compared to congruent conditions,^92^ suggesting misaligned feedback actively disrupts the neurofeedback learning process.^90,93,94^ If the goal is to induce neural plasticity through neurofeedback, misleading signals could reinforce maladaptive brain patterns.^95^ Thus, careful consideration of control conditions that do not interfere with the neurofeedback process is crucial for clinical research, especially among vulnerable populations.^96^

## Limitations

Several limitations warrant consideration. One issue is the use of different MRI scanners, as hardware variability can impact BOLD signal estimates. However, we mitigated this through within-subject longitudinal measurements on the same scanner and balanced cross-condition comparisons across locations.^97^ Another limitation is the absence of double-blind design. Additionally, without a mental-noting-only control condition, we cannot isolate neurofeedback-specific effects. While these findings replicate previous results, larger sample sizes are needed for verification. Finally, a single neurofeedback session targeting the superior temporal gyrus may be insufficient for learning cognitive strategies, suggesting the need to study how session parameters influence outcomes.

## Conclusion

This study demonstrates that real-time fMRI neurofeedback combined with mental- noting practice can effectively modulate brain activity and connectivity patterns implicated in AHs. The observed reduction in functional connectivity between secondary AC and rDLPFC in the Real-NFB group, along with its trend-level relationship with AHs reduction, provides preliminary evidence for the therapeutic potential of this intervention. These results highlight the promise of combining targeted neurofeedback with mindfulness techniques as a novel therapeutic approach for AHs, while also emphasizing the importance of appropriate control conditions in clinical neurofeedback research. Future directions include testing a double-blind study design to minimize bias and using larger sample sizes to replicate findings. Finally, future studies should investigate how the duration and frequency of neurofeedback sessions influence neural activity, connectivity, and symptom severity.

## Acknowledgments

This work was supported by the National Institute of Health (Grant No. R61MH113751 to MAN and SWG).

CCCB, KO, AKS, MAN and SWG designed research; CCCB, JZ, FM, YL, LS, AA, MH, AKS, SWG and MAN performed research, and analyzed data; JH performed data curation and analytical support; CCCB, JZ, FM, AKS, MAN and SWG wrote the manuscript. All authors read and approved the manuscript. We thank all the participants who took part in our study and the staff at Massachusetts Institute Technology, Northeastern University Biomedical Imaging Center, McLean Hospital, and Veterans Affairs for assistance with data collection.

