## supplemental for "Real-time fMRI neurofeedback modulates auditory cortex activity and connectivity in schizophrenia patients with auditory hallucinations: A controlled study"

**Supplementary Materials**

Figure S1. CONSORT diagram.	1

Figure S2. Replication of Okano et al. 2020 results: Primary auditory cortex

activation changes post-NFB**.**	4

Figure S3. Whole brain analysis of functional connectivity changes of primary

auditory cortex post-NFB	6

Table S1. MRI Scanning and Study Design Information	9

Table S2. Mean and standard deviation values for activation and functional

connectivity pre- and post-intervention.	10

Replication Analysis, Results & Discussion	3

Mindfulness Training Instructions	11

MRI Acquisition	15

References	16

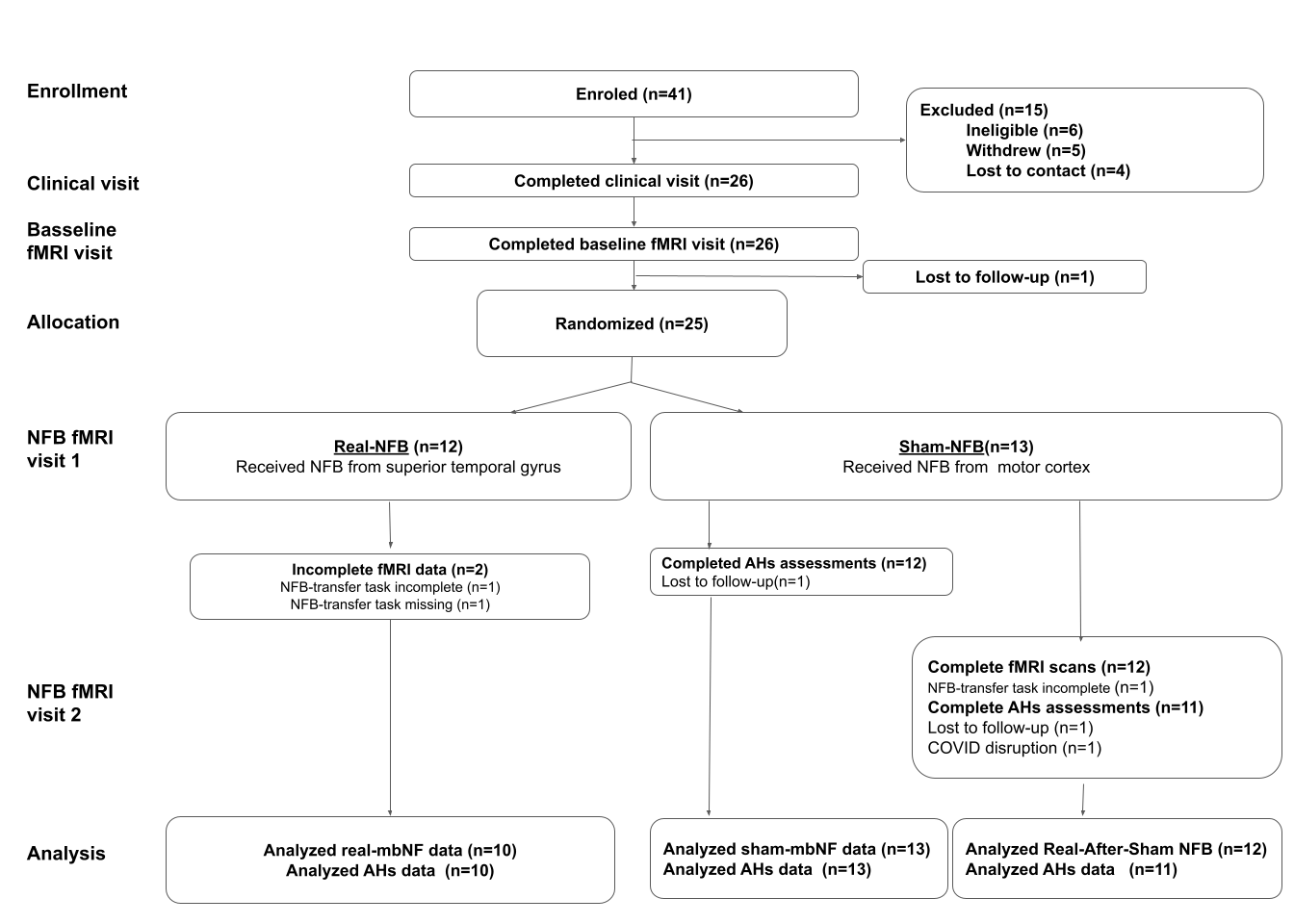

##### Figure S1. Consolidated Standard of Reporting Trials (CONSORT) diagram.^1^ Note that the CONSORT diagram relates to data presented in the current paper, which is a subset of a multi-visit study. Information of the full study can be found on clinicaltrials.gov (NCT03504579) and in other publications (Zhang et al., Submitted; Morfini et al.^2^) AH: auditory verbal hallucinations; NFB: neurofeedback; rsfMRI: resting state functional magnetic resonance imaging.

### *Replication Results*

##### ***Primary auditory cortex activation***

We tested whether activation in the right PAC-ROI (sphere of 10 mm around the peak coordinates (MNI X,Y,Z=60, -18, 10) reported in our previous study)^16^ was changed by NFB training using a two-way ANOVA. There was a significant effect of Time F(2, 20)=2.41, *p*=.02, Cohen's *f^2^*=.30 (MNI X,Y,Z=54, -22, 8, non-parametrically SVC at *p*<.001; **Figure S2A**), with no effect of Group and no Time X Group for Real-NFB and Sham-NFB. This was confirmed by post hoc paired t-tests within the conditions showing significant difference from pre- to post-intervention for Real-NFB [*t*(9)=5.12, *p*=.0003, Cohen’s *d*=1.26, **Figure S2B**] and for Sham-NFB [*t*(12)=2.48, *p*=.02, Cohen’s *d*=.82 **Figure S2B**]. Additionally, when assessing the Real-after-Sham-NFB and Sham-NFB conditions, there were no significant Time, Condition or Time x Condition effects. However, paired t-tests within the conditions showed reduction in PAC activation from pre- to post-intervention [*t*(10)=2.0, *p*=.03, Cohen’s *d*=.92, **Figure S2C**]. This reflects an overall reduction in PAC activation on the first Sham visit, with additional reduction in PAC activation seen for the second visit in the Real-after-Sham-NFB condition (see mean and standard deviation values for pre- and post-intervention in **Table S2)**. We also assessed the relationship between baseline AHs scores and activation in the right PAC-ROI. We found a significant positive relationship between baseline AHs scores and pre-intervention right PAC activation during other-voice blocks in all subjects (MNI X,Y,Z=60, -26, 13, BA41, n=23, r=.71, non-parametrically SVC at *p*<.001; **Figure S2D & E**).

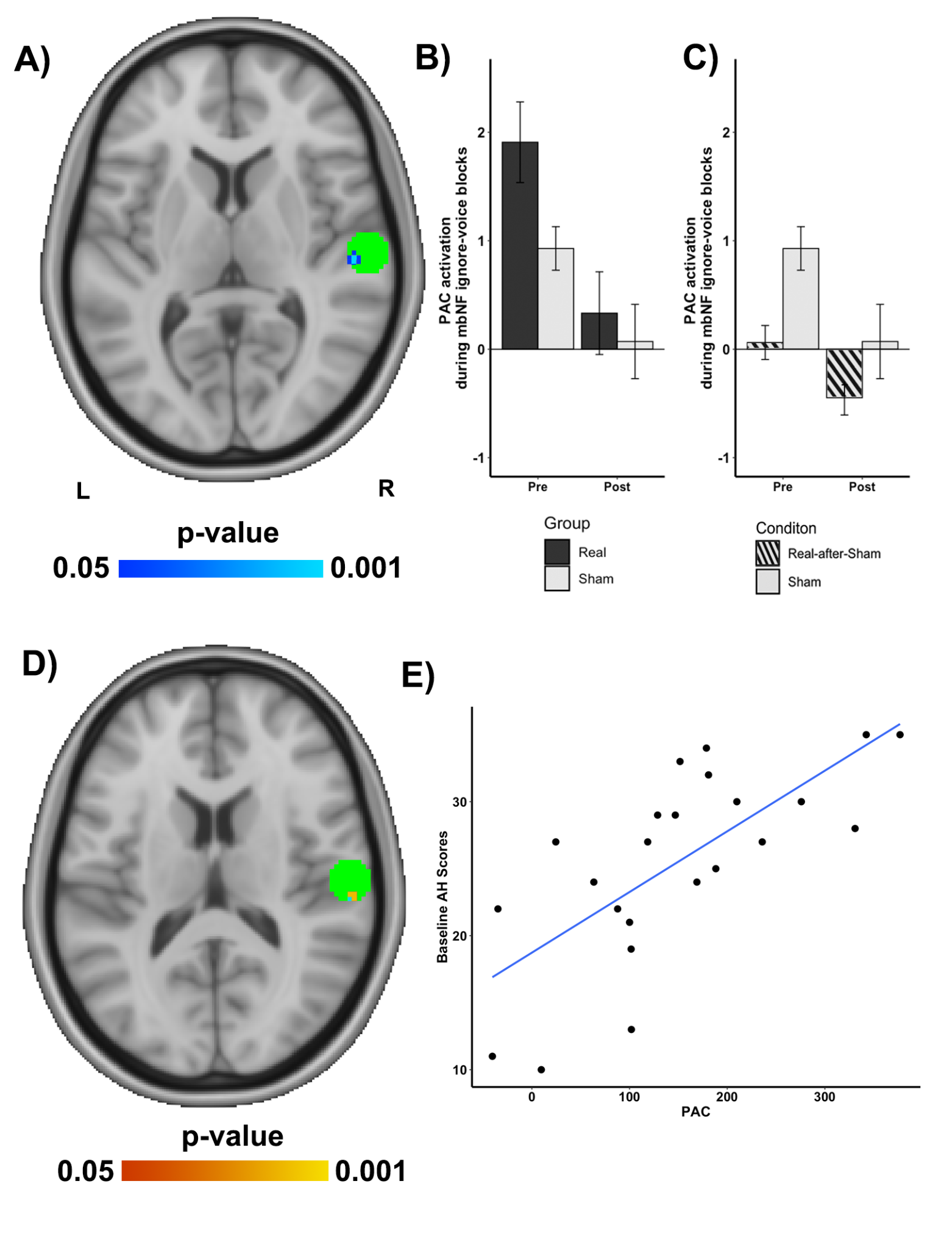

#### **Figure S2. Replication of Okano et al. 2020 results: Primary auditory cortex activation changes post-NFB.** (A) Axial sections of the brain depicting an a priori primary auditory cortex (PAC) ROI sphere of 10 mm around the peak coordinates (MNI X,Y,Z=60, -18, 10, BA41, green sphere) reported in Okano et al., 2020.^3^ Axial section of the brain depicting voxels in the PAC where activation reflects a significant Time effect for all conditions. (B) Significantly decreased PAC activation pre-to-post intervention in the both Real- and Sham-NFB conditions. (C) Additional reduction in PAC from pre-to-post intervention in the Real-After-Sham-NFB condition relative to Sham-NFB. (D) Axial section of the brain depicting voxels in the PAC in which baseline activation during ignore-voice blocks (E) correlated significantly and positively with AHs scores before the intervention. All bars reflect mean and all error lines reflect standard error of the mean; statistics are nonparametric and FWE-pTFCE small volume corrected.

##### ***Effect of NFB on functional connectivity during ignore-voice***

***Primary Auditory Cortex Connectivity***

We tested whether functional connectivity between PAC and the rest of the brain was changed by NFB. Standard psychophysiological interactions (gPPI) analysis was used from the PAC cluster found within the Okano et al., 2020 ROI from the two-way ANOVA between Real-NFB and Sham-NFB groups (see blue cluster peak: 54,  -22,  8; BA41 within green ROI in **Figure S2A**) and whole brain analysis during the *ignore-voice* contrast in a second two-way ANOVA. During the *ignore-voice* blocks, there was a significant effect of Time [*F*(2, 20)=2.31, *p*=.03, Cohen's *f^2^*=.43 (rDLPFC, *n*=22: *X*=38, *Y*=28, *Z*=24; BA9; non-parametrically at *p*<.001; **Figure S3A**], with no effect of Group and no Time X Group interaction. This was confirmed by post hoc paired t-tests within the groups showing significant difference from pre- to post-intervention for Real-NFB [*t*(10)=1.93, *p*=.04, Cohen’s *d*=0.86, **Figure S3B**] and for Sham-NFB [*t*(10)=1.87, *p*=.04, Cohen’s *d*=.86 **Figure S3B**]. Additionally, when assessing the Real-after-Sham-NFB and Sham-NFB conditions, there were no Time, Condition, or Time x Condition effects. Paired t-tests within the conditions showed reduction from pre- to post-intervention (t(10)=2.0, *p*=.03, Cohen’s *d*=1.11, **Figure S3C**). These effects reflect an overall reduction in PAC-rDLPFC functional connectivity across conditions on the first visit, with additional reduction seen for the second visit in the Real-after-Sham-NFB condition (See mean and standard deviation values for Pre- and Post-intervention in **Table S2**).

**Figure S3. Whole brain analysis of functional connectivity changes of primary auditory cortex post-NFB.** (A) Axial section of the brain depicting voxels in the PAC and right dorsolateral prefrontal cortex (rDLPFC) where functional connectivity reflects a significant Time effect. (B) Significantly decreased PAC-rDLPFC functional connectivity pre-to-post intervention in the Real-NFB and Sham-NFB groups. (C) Significantly decreased activation pre-to-post intervention in the Real-after-Sham-NFB and Sham-NFB conditions. All bars reflect mean and all error lines reflect standard error of the mean; statistics are nonparametric and FWE-pTFCE small volume corrected. PAC: Primary Auditory Cortex.
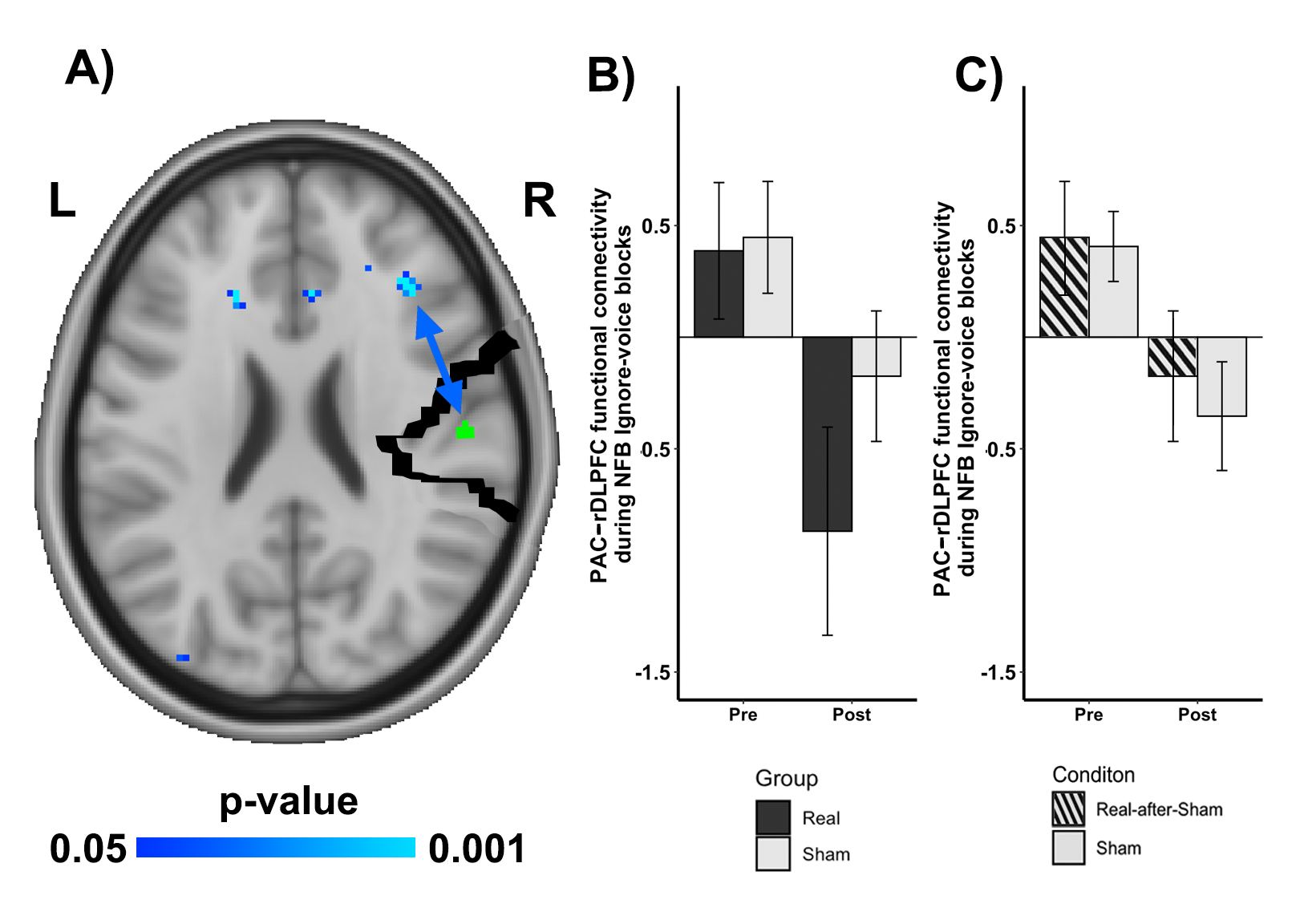

### *Effect of NFB on activation in cognitive control regions during ignore-voice*

When assessing the activation change for the rDLPFC obtained during gPPI analyzes during the *ignore-voice* blocks, there was no significant Time X Group interaction (*p*>.05, Cohen’s *f^2^*<.05), such that activation for the rDLPFC during *ignore-voice* was maintained after receiving both Real-NFB or Sham-NFB.

### Discussion

Our study aimed to validate and extend the findings of our previous research on NFB training aimed at reduction of AHs in schizophrenia.^3^ The results provide several important insights into the effects of NFB on PAC activation and functional connectivity, as well as the relationship between AHs symptoms and neural patterns.

##### ***Reduction in Primary Auditory Cortex (PAC) Activation***

The significant effect of Time on right PAC activation across both Real-NFB and Sham-NFB conditions suggests that the intervention influenced auditory processing regardless of NFB nature. The reduction in PAC activation from Pre- to Post-intervention in both conditions indicates a general effect of the experimental procedure. This could be due to non-specific aspects related to reallocation of brain resources during the mental-noting practice^4,5^ as well as more generally attributed to habituation,^6^ learning effects.^7,8^ Of note, the Real-after-Sham-NFB condition showed additional reduction in PAC activation during the second visit. This suggests a potential cumulative effect of the intervention even when the initial sessions were sham.^9,10^ This finding highlights the possible benefits of meditation enhanced interventions in the treatment of AHs as well as the complexity of NFB effects and the potential importance of extended training periods. The positive correlation between baseline AHs and pre-intervention PAC activation during 'other voice' blocks supports the hypothesis that heightened auditory cortex activity is associated with more severe hallucinations. This replicates previous findings^3^ and strengthens the rationale for targeting PAC activity in AHs treatment.^11^ This is also in line with studies that have shown that patients with schizophrenia and AHs often exhibit abnormal activation patterns in the PAC during auditory processing tasks.^12,13^ This dysregulation of the PAC is thought to contribute to the perceptual quality of AHs, making them subjectively vivid and similar to hearing actual voices.^13,14^ Interestingly, neurofeedback training has demonstrated the ability to modulate early auditory processing in the PAC.^15^ This modulation of PAC activity through neurofeedback has been associated with a reduction in AHs symptoms, suggesting a potential therapeutic mechanism.^16,17^ These findings highlight the involvement of the PAC in the neural mechanisms underlying AHs additional to the previously discussed secondary AC involvement and suggest that targeted modulation of early auditory processing through neurofeedback may also be a promising approach for reducing AHs in schizophrenia and in other disorders.

##### ***Reduced Functional Connectivity between PAC and*** rDLPFC

The observed changes in functional connectivity between PAC and right dorsolateral prefrontal cortex (rDLPFC) during 'ignore-voice' blocks are particularly intriguing. The reduction in PAC-rDLPFC connectivity across both Real-NFB and Sham-NFB conditions, with additional reduction in the Real-after-Sham-NFB condition, parallels the patterns seen in PAC activation. This decrease in connectivity could reflect engagement of cognitive control brain areas and reallocation of attentional resources away from auditory processing and towards other bodily sensations during mental-noting practice.^18,19^ Accordingly, this change in connectivity could be driving the effects of reduced PAC activation discussed in the previous section.

##### Maintenance of rDLPFC Activation

The maintenance of rDLPFC activation during 'ignore-voice' blocks, despite decreased connectivity with PAC, is noteworthy. This suggests that while the communication between auditory and executive control regions may be reduced due to reallocation of attentional resources,^18,19^ the engagement of cognitive control processes remains stable.^20,21^ This indicates that regardless of condition assignment, participants were properly engaging in mental-noting, hence exerting cognitive effort with reallocation of attention away from auditory stimuli and suggests that the intervention influenced both auditory and cognitive control processing regardless of NFB nature.

In conclusion, our replication study supports the potential of NFB as an intervention for AHs in schizophrenia, while also highlighting the complexity of its effects. The observed changes in PAC activation and functional connectivity provide a foundation for understanding the neural mechanisms underlying NFB efficacy. However, the similarities between Real-NFB and Sham-NFB effects underscore the need for careful experimental design and consideration of non-specific factors in future research and clinical applications.

**Supplementary Tables**

Table S1. MRI Scanning and Study Design Information.

|  | Real-NFB  (n=10) | Sham-NFB  (n=13) | t \| χ2 | p | sig |
| --- | --- | --- | --- | --- | --- |
| **MRI scanner (n)** |  |  |  |  |  |
| Site | NEU (6)  MIT (4) | NEU (8)  MIT (5) | ~0 | 1 | n.s. |
| Type | Prisma (8)  Trio (2) | Prisma (12)  Trio (1) | ~0 | 1 | n.s. |
| Head coil channels | 64 (8)  32 (2) | 64 (12)  32 (1) | ~0 | 1 | n.s. |
| **Individualized masks** |  |  |  |  |  |
| STG size (n voxels) | 36.3 (±26.12) | 43.37 (±33.55) |  |  |  |
| STG laterality | Right (10) | Right (13) |  |  |  |
| SMC size (n voxels) | 1652.5 (±227.63) | 1799 (±331.24) |  |  |  |
| SMC laterality | Bilateral (10) | Bilateral (13) |  |  |  |
| **Time between visits (days**) |  |  |  |  |  |
| Screening and Baseline | 19.38 (±8.94) | 25.9 (±15.59) | 1.05 | 0.31 | n.s. |

MIT, Massachusetts Institute of Technology; NEU, Northeastern University; NFB, real-time fMRI neurofeedback; fMRI, functional magnetic resonance imaging; n.s., not significant at a p-value < 0.05; sig, significance.

Table S2. Mean and standard deviation values for activation and functional connectivity pre- and post-intervention.

|  | Real-NFB  (n=10) | | Sham-NFB  (n=13) | | Real-after-Sham-NFB  (n=11) | |
| --- | --- | --- | --- | --- | --- | --- |
|  | pre | post | pre | post | pre | post |
| AHs-score | 25.90 (7.09) | 20.00 (7.13) | 25.23 (7.48) | 22.33 (7.31) | 21.90 (7.5) | 21.00 (10.66) |
| Motion | 9.30 (4.5) | 9.10 (6.43) | 7.15 (3.78) | 8.23 (4.26) | 5.90 (3.38) | 8.45 (5.80) |
| Activation β |  |  |  |  |  |  |
| SAC (BA 22) | 1.40 (1.49) | 2.7 (1.25) | 0.72 (0.68) | 0.95 (1.20) | 0.5 (1.08) | 0.06 (0.66) |
| PAC (BA 41) | 1.90 (1.17) | 0.33 (1.20) | 0.92 (.72) | 0.07 (1.23) | 0.06 (0.51) | -0.44(0.52) |
| BA 10 | 1.26 (2.50) | 0.63 (2.75) | 1.64 (1.30) | -0.21 (1.45) | 0.56 (1.48) | 0.80 (1.29) |
| Connectivity z-score |  |  |  |  |  |  |
| SAC & BA 10 | 0.60 (1.16) | -0.61(1.25) | 0.60 (1.38) | 1.14 (1.14) | -0.39 (1.16) | -0.17(1.07) |
| PAC & BA 10 | 0.38 (0.96) | -0.86 (1.47) | 0.44 (0.86) | -0.17 (1.01) | 0.40 (0.51) | -0.35 (0.80) |

AHs: auditory visual hallucinations; PAC: Primary auditory cortex; SAC: secondary auditory cortex; BA: Brodmann area; NFB, real-time fMRI neurofeedback.

### Mindfulness Training

### Mental Noting Practice

During the neurofeedback visit, the experimenter followed the following scripts to guide the participants through the mental noting practice.

**Part 0: Short story test - baseline**

*“Before we teach you anything. Let’s do a simple memory test. You’ll hear a story and let’s see how many details you can remember from it.*

*Next Saturday is Mark's birthday. Mark's friends are going to hold a secret party for him. Tim is one of Mark's best friends and loves to prank other people. Although everyone usually eats a sweet and well-decorated cake on their birthday, Tim is going to make one that is salty. He can’t wait to have Mark try the cake and laugh at his response!”*

**Part 1: Explanation**

*“In today’s experiments, we will be studying a special type of attention called effortless awareness. An example of effortless awareness is to become aware that hearing the sound of my voice does not require any special effort on your part. In contrast, when you hear sounds and immediately try to understand or make sense of them, you experience an effortful state.*

*Does that make sense?*

*The same effortful state is, for example, if you were in an argument recently, it is easy to get caught up in emotions by reliving the situation in your head. The thoughts or situation aren’t the issue, but how you are relating to them – being caught up- in them.*

*Does that make sense?*

*Today we would like you to practice a simple attention technique that will allow you to rest in effortless awareness while ignoring all sounds. This technique is called the noting practice. During noting practice you simply note what is most predominant in your experience from moment to moment. That is, you just become aware of your senses, like seeing, feeling, hearing, and thinking. Just note silently to yourself whatever is at the forefront of your awareness at any moment. For example, I might note seeing, because I’m reading this, and then I notice my shoulders are tense so I note feeling, and then I start thinking that my shoulders are tense so I note thinking. The sequence would go like this: seeing, seeing, seeing, feeling, feeling, thinking, thinking and so on. Whatever is most predominant in your sensory awareness from moment to moment, just note it. You don’t need to describe specifically what you’re experiencing, but we’d like you to note which senses you are using and let it go after you have noted it.*

*Does that make sense?*

*At the beginning it is recommended to pace the rhythm to once per second. But it’s up to you.*

*I will provide an example (15s, note approx.. once per second)*

*Notice sometimes I dwell on ‘hearing’ or ‘thinking’ for more than 2 seconds. In those cases, I use an ‘anchor’ so that I force myself to switch my attention to something other than hearing or thinking. For example, I use my right big toe as an anchor. That means whenever I notice I say ‘hearing’ or ‘thinking’ for 2 or more times, I force myself to pay attention to how my big toe is feeling. It provides an easy ‘out’.*

*Does that make sense?*

*So let’s try to help you pick an anchor. A body part can be an option. What would you like to use as your anchor?”*

**Part 2. Participant practices noting without external interference.**

“*Could you try this now, just noting seeing, feeling, smelling, tasting, etc., just noting anything that is at the forefront of your attention for the next 10 seconds or so? And could you note out loud this time so I can follow you? I will stop you in 10 seconds.*

*Don’t worry about doing it perfectly; just do the best you can. If your mind wanders or you get caught up or swept away by something, no problem; when you become aware again, just start again by noting whatever is most predominant in your awareness. “*

**Part 3. The participant practices mental noting while the experimenter tells a short story.**

*“Great. Now let’s practice noting with some background sound. This time, after I say ‘start’, you will start practicing noting as before - you can choose to say it out loud or note in your head without speaking out loud. But at the same time, a story will be played. Your goal is simply to keep going on with the noting practice even while the story is playing .. I will say ‘ready, go’ and you’ll have 5 seconds before the story starts. When the story ends, you’ll hear ‘stop’. Are you ready?”*

**Audio stories that participants heard during mental noting**

Audio files for the current project can be found at:

<https://github.com/cccbauer/MindfulnessTraining/tree/main>

Audio Story 1

1. Cathy is going on a field trip to a dairy farm. When Cathy gets to the farm, the farmer shows her how he makes fresh milk and creamy cheese. At the end of the day, the farmer said everyone can bring something home as a present for their parents. Cathy's mom and dad love pizza and Cathy's favorite food is macaroni and cheese. She knows exactly what to bring.

Audio Story 2

2. Alex the bear lives on the mountain. One sunny day, Alex was out looking for food. He saw a beehive on a tall tree. Alex climbed up and stuck his paw inside the beehive. He took his paw out and ate a lot of honey! Alex wanted to take a break, so he went to sit down in the shade under a big tree. Alex closed his eyes.

Audio Story 3

3. Zoey is going on vacation with her family by the ocean. She is so excited! She packed her favorite pair of sunglasses and her favorite bathing suit. They drive to the coast. But they see a lot of clouds in the sky and everyone’s hair is blowing in the strong wind. Soon rain starts pouring. Zoey is a little sad.

Audio Story 4

4. Grandpa owns a garden. He works there all year round. In March, he uses the shovel to dig holes in the soil and plants the potato seeds. From March to October, Grandpa waters and fertilizes the plants. In October, Grandpa carefully digs out the potatoes. In the winter, Grandpa takes a break.

*“If noting practice is done well, you will notice that whatever you hear, you may not understand as well - the content ‘fades away’. That’s the goal for the noting practice. So let’s try next how you can do this.*

*Remember you can use your anchor if you notice yourself getting involved in the story.”*

##### ***MRI Data Acquisition***

MRI data were acquired with 3T MRI scanners, either at Massachusetts Institute of Technology with a Trio (n=5) or a Prisma system (n=5) , or at Northeastern University (NEU, n=13) with a Prisma system. Each participant underwent MRI scanning sessions at the same site.

*Trio MR System:* All MRI data for 5 participants were performed using a 32-channel head coil. Anatomical data were acquired using a T1-weighted magnetization-prepared rapid acquisition gradient-echo (MPRAGE) pulse sequence: resolution=1 mm isotropic, repetition time (TR)=2,530 ms, echo time (TE)=1.61 ms, flip angle (FA)=7°, and inversion time (TI)=1,200 ms. For all functional data, the blood oxygen level dependent (BOLD) signal was measured using a gradient-echo, echo-planar imaging pulse sequence (EPI): resolution=3.5mm isotropic, TR=2,000 ms, TE=30 ms, FA=90°. For each task, two runs with opposite phase encoding direction (anterior-posterior (AP-PA)) were acquired.

*SIEMENS MAGNETOM Prisma:* All MRI data for 17 participants were performed using a 32-channel (n=4, MIT) or 64-channel head coil (n=13, NEU). Anatomical data were acquired using a T1-w MPRAGE: resolution=1mm (MIT) or 0.8mm (NEU) isotropic, TR=2,530 ms, TE=1.7 ms, FA=7°, TI=1.4. For all functional data, the BOLD signal was measured using EPI sequence with imaging parameters: resolution=2 mm isotropic, TR=1.2 s, TE=30 ms, FA=72°.
